# Genetic incompatibility combined with female-lethality is effective and robust in simulations of *Aedes aegypti* population control

**DOI:** 10.1101/316406

**Authors:** Maciej Maselko, Stephen Heinsch, Siba Das, Michael J. Smanski

## Abstract

Recent reports of CRISPR/Cas9-based suppression gene drives in insects underscore the challenge of overcoming genetic resistance. Here we present results from agent-based simulation modeling of a novel Field-Amplified Male Sterility System (FAMSS) that outperforms suppression gene drives when challenged with genetic resistance. FAMSS combines a recently described synthetic genetic incompatibility approach with previously demonstrated female-lethality constructs. Our results suggest that FAMSS will be an effective strategy for temporally and spatially self-limited suppression of the disease vectoring mosquito, *Aedes aegypti*.

## Introduction

Genetically engineered (GE) biocontrol agents can provide effective, species-specific population suppression of disease vectors^1–3^, agricultural pests^4,5^, and invasive species^6,7^. Several different GE biocontrol strategies have been proposed, each with unique strengths and weaknesses^8^. Key differences exist in the mechanisms resulting in different degrees of scalability (e.g. as a result of egg versus adult release) and differences in the likeliness and fitness of resistance mutations.

Release of insects carrying dominant lethals (RIDL) is a proven technology that has been demonstrated in various insects^9,10^. In RIDL, a ‘self-limiting’ gene is controlled by a small-molecule repressible promoter. The self-limiting gene is one that reduces the fitness of the organisms upon expression, either by causing lethality or a deleterious phenotype. Propagation of the GE biocontrol agent in the presence of the small-molecule repressor keeps the self-limiting gene turned off. Hybrids between released RIDL adults and wild mates experience embryonic or late-onset lethality in the absence of the repressor due to the unregulated expression of the self-limiting gene.

Sex-ratio biasing GE biocontrol agents have been engineered via several mechanisms^6,11^. Female-lethal strains can be designed analogous to RIDL insects, but using a repressible female-specific promoter or alternative splicing variant to control expression of the self-limiting gene. Alternatively, sex-ratio biasing can be engineered with X-shredder designs that express nucleases to cleave the X-chromosome during meiosis, resulting in only male offspring^12,13^. In species whose sex determination is influenced by both genetics and environment, YY females can be produced for an approach known as ‘Trojan Y chromosomes’^14^. Released YY females have only male offspring and skew the sex ratio in subsequent generations.

The development of sequence-programmable nucleases, especially CRISPR/Cas9-based systems, has accelerated the construction of threshold-independent meiotic gene drives^12,15,16^. In meiotic gene drives, ‘drive’ alleles encode a sequence programmable nuclease expressed during meiosis^17^. Expression of the nuclease leads to cutting of a wild allele and replacement with the drive by gene conversion via homology-directed repair. Using promoters that direct expression of the drive allele early in gametogenesis, an organism heterozygous for the drive allele in somatic tissue will become homozygous in the germline, only passing on the drive allele to its offspring. ‘Suppression gene drives’ (SGDs) target a haplosufficient essential gene and will cause a population to crash once the drive allele has spread through the population^12^.

We have recently demonstrated a strategy for GE biocontrol called synthetic genetic incompatibility (SGI)^18^. SGI approximates the behavior of a single-locus extreme underdominance system, which have been noted for their theoretical applications in population control^19^. SGI is engineered by designing programmable transcription activators (PTAs) to drive lethal overexpression or ectopic expression of endogenous genes in hybrid embryos that are produced when GE organisms mate with wild-type. Thus, any mating event between and SGI organism and wild-type will fail to produce viable offspring^18^. A variation of SGI which specifically targets gametogenesis by limiting PTA expression to the germline would result in hybrids that are viable but sterile.

Wide-spread use of GE insects for population control is likely in the near future. Technologies that use RIDL for the control of agricultural pests^4^ and disease vectors^20^ have been approved for field trials in the US by the USDA, FDA, and EPA. Sex-ratio biasing constructs have been developed in several insect species as well as in vertebrate pests^6, 21–23^. Meiotic gene drives have been constructed in laboratories and tested in caged studies to suppress populations of mosquitoes and fruit flies^12,24,25^. All of the components required to engineer an SGI strain have been shown to function properly in D. melanogaster^26^. Predicting the performance qualities of these population control strategies using computational models and simulations is an important prerequisite to laboratory or field trials.

Here we introduce a new GE biocontrol strategy, named Field-Amplified Male Sterility System (FAMSS) that combines sterilizing SGI with female-lethality. We assess the performance of FAMSS and alternative GE biocontrol agents using a spatially-defined agent-based simulation model of *Aedes aegypti* populations. As the primary vector of multiple viruses including Dengue, Zika, and Chikungunya, *Ae. aegypti* is a critical target for applications in disease control^27^. We show that FAMSS is a spatially and temporally self-limiting strategy to control *Ae. aegypti* populations. Lastly, we show that the FAMSS strategy outperforms suppression gene drives in the context of genetic resistance because of differences in relative fitness of resistant organisms.

## Results

### Agent-based simulation model for Aedes aegypti

We developed an agent-based model for *Aedes aegypti* that allows us to track the genotype of every individual in an arbitrary number of neighboring populations (Figure 1 and Supplementary Note 1). The model is an adaptation of the simulation described by Dye^28^ and is based on empirical data from the 1966 World Health Organization Aedes Research Project at the half hectare Wat Samphaya in Bangkok, Thailand^29^. Whereas the Dye model treats adulthood as a single black-box, our model includes more detailed progression through multiple gonotrophic cycles (GCs). Because of this, we incorporate additional empirically-defined parameters on daily survivorship, egg deposition per GC, and gestation time^30^. Unlike more recent simulation models that track fine-scale movements of mosquitoes between specific habitat features such as standing water pools, and blood sources^31,32^, our model only considers spatial movements in terms of migration between neighboring populations. This last feature allows us to predict what will happen in border regions where GE mosquitoes can interact and crossbreed with non-targeted wild-type populations.

**Figure 1.**
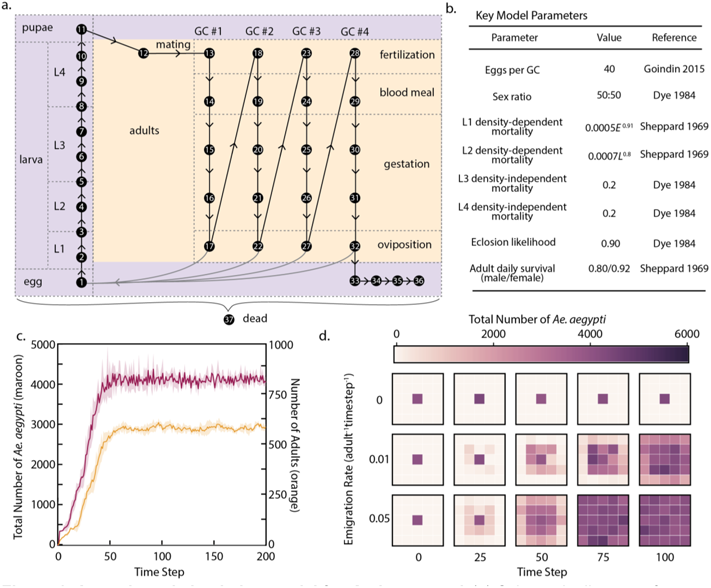
Agent-based simulation model for *Aedes aegypti*. (a) Schematic diagram of agent-based model, showing all possible life-stages (black nodes) for female mosquitoes. GC = gonotrophic cycle; L1-4 = larval instar stages I-IV. (b) List of key model parameters. (c) Results from simulation of a single population with initial size of 500 individuals (randomly distributed across life stages). Maroon trace shows mean number of agents with lifestage between 2 and 36. Shadow shows standard deviation from five independent replicates. Orange trace shows mean number of adults with shadow denoting standard deviation from five independent replicates. (d) Migration between neighboring populations as a function of time. Results from three experiments, each run in triplicate, showing migration of *Ae. aegypti* from a seed population (center square on grid) to surrounding populations. Square color represents total number of individuals (lifestage 2-36) in each population at the time step labeled below.

Within a single population, individual agents in our model progress through a series of 37 discrete states (life-stages) in time-steps of 1.57 days (Figure 1a,b). These states include larval, pupal, and adult stages. Adult females mate with a random male in the population during early adulthood. Sperm from this mating event is stored and used to fertilize eggs at the start of up to four gonotrophic cycles (GCs). However, the daily mortality rates are sufficiently high so that the average female will only go through one GC. For mating events that produce embryonic lethality, inviable embryos are ‘killed’ between life-stages 1 and 2 (hatching of eggs) (Figure 1a). Specific examples of mating events that result in embryonic lethality are described in subsequent sections.

A wild-type population of mosquitoes reaches equilibrium due do density-dependent mortality rates during L1 and L2 instars. We have tuned the larval density-dependent mortality rates within the range of measured values such that the steady state population size fits observations from the Bangkok study^29^ (Supplementary Note 2). In our model, equilibrium is reached at approximately 4000 total agents including 600 adults (Figure 1c). The number of adults estimated in the Bangkok study ranges between 460 and 2130 during different months^29^. Further, our simulation model shows expected behavior characteristics for *Ae. aegypti* populations including monotonic stability and robustness to changes in fecundity^28^. We have included a rough spatial component in our model that allows tracking of allele frequencies between neighboring populations. We include a parameter termed ‘migration rate’ that describes the likelihood that a given adult mosquito will migrate into a neighboring population at each time step (Figure 1d). Studies on the migration rates of individual mosquitoes in the modeled Wat Samphaya population show that movement is limited to only tens meters per day^29^, which coincides with the range of migration rates tested here. All of the parameters in our model can be changed to account for the unique population dynamics specific to a location of interest.

### Efficacy of genetic population control methods on individual populations

We performed 200 time step simulations for alternative GE population control methods, including female-lethality, SGI, and SGD (Figure 2a-c). For each strategy, we simulated a one-time release of GE mosquitoes at different starting frequencies of GE and wild-type. We performed each experiment in triplicate and report the mean population composition at each time-step as well as the standard deviation of the mean. Our results confirm the expected behavior of each modeled control strategy.

**Figure 2.**
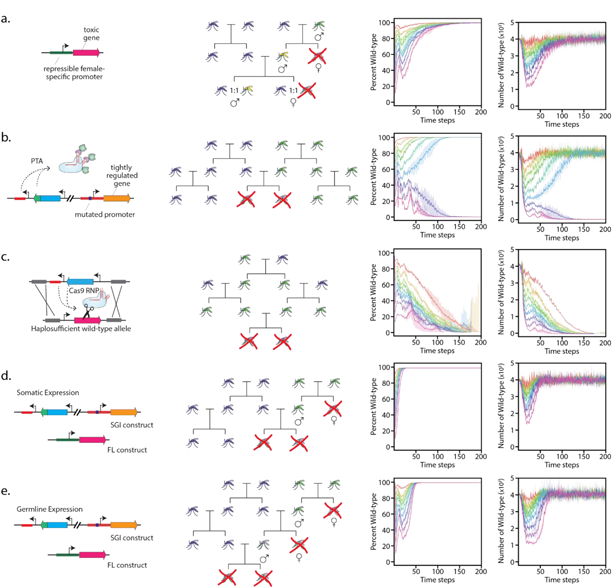
Comparison of alternative genetic population control methods in one-time release. Schematic drawing of genotype (left), mating outcomes of wild-type (purple) and GE (green) mosquitoes (center), and modeled performance at different release rates (right) for (a) flightless female (FL), (b) synthetic genetic incompatibility (SGI), (c) homing suppression gene drives (SGD), (d) self-stocking incompatible males (SSIMS), and (e) field amplified male sterility system (FAMSS). Performance graphs show the fraction simulated population comprised of GE mosquitoes (left plot) and the total number of wild-type mosquitoes in the simulation (right plot). Each simulation starts with steady-state (4000) wild-type agents and enough GE agents to comprise 10% (red), 20% (orange), 30% (light green), 40% (dark green), 50% (light blue), 60% (dark blue), 70% (indigo), 80% (violet), or 90% (magenta) of the starting population. Lines correspond to mean from three independent experiments, while back-shading denotes standard deviation. PTA, programmable transcription activator; RNP, ribonucleoprotein.

We model the FL system as the previously reported ‘flightless female’ *Ae. aegypti* ^33–35^. Released GE mosquitoes are given the genotype ‘LL’ (wild-type are ‘ll’), and the flightless phenotype is present in females with at least one ‘L’ allele. We simulate the flightless female phenotype by removing them from the model at the first adult life-stage. The FL strategy displayed a temporally self-limiting population suppression that scaled with initial release rate (Figure 2a). After initial release, the fraction of the population comprising wild-type individual eventually returned to 100% as the FL genotype was diluted out.

The SGI release strategy was modeled for population replacement applications where both male and female SGI mosquitoes are released. We encoded the SGI genotype as two independently segregating genetic loci. ‘P’ denotes a mutated promoter in a lethal over-expression gene target and ‘T’ denotes the presence of a programmable transcription activator ^18,26^. GE mosquitoes have the genotype ‘PPTT’, and wild-type have the genotype ‘pptt’. Any embryo that contains at least one ‘T’ allele and at least one ‘p’ allele is inviable. With release of both sexes, SGI is expected to act as a threshold-dependent gene drive. At frequencies greater than 50%, the SGI genotype is expected to drive to fixation. At frequencies less than 50%, the wild-type genotype is expected to drive to fixation. We observe a clear threshold of 50% population composition in the simulation models (Figure 2b), but note that achieving fixation required an initial release of 70% SGI agents. The reason for the discrepancy has to do with differing life-stages of the initial populations of wild-type versus engineered individuals. In our simulations, the starting wild-type population contains a distribution of life stages (*e.g*. larval, pupal, adult) that represents a steady-state untreated population. The released bolus of GE organisms only contains individuals in the egg/larval stages. We do this to simulate realistic biocontrol applications. Because of density dependent mortality rates at larval stages, an initial release rate of 50% is quickly reduced to a sub-50% population frequency (Figure 2b). It is noteworthy that even for some release rates that do not result in fixation of the SGI genotype, there is a substantial wild-type population suppression effect that lasts 50-100 time steps (Figure 2b, right).

We modeled SGD applications using a genotype similar to that previously developed suppressive meiotic drive in *Ae. aegypti*^12,36^. Released SGD mosquitoes have the genotype ‘GW’ (compared to ‘WW’ for wild-type mosquitoes), signifying that they are heterozygous for the drive allele in somatic tissue. For the purpose of the simulation in Figure 2c, we assumed ideal drive behavior (100% homing rate, 0% non-homologous end joining repair) to show that this threshold-independent gene drive functions as predicted in our simulation model. At release rates between 90% and 10%, the SGD always drives to fixation. In all cases, the population of wild-type mosquitoes is completely suppressed with only the single initial release of GE biocontrol agents. (Figure 2c right).

### Combining genetic incompatibility with female lethality for genetically-encoded sterile male release

Next, we simulated a combination GE control method that entails engineering both FL and SGI into a single biocontrol strain. We named this approach Self-Sorting Incompatible Male System (SSIMS), because GE males and females can be released together to suppress a wild population. There are two feasible release strategies. First, the released individuals could be reared in the presence of tetracycline (to repress FL construct) and released as adults. This scenario would be ideal for pests or organisms that live and reproduce for many years (*e.g*. invasive carp), as SSIMS males and females will both suppress the population when they mate with wild-type. If they mate with each other, only SSIMS males are viable and will amplify the suppressive effect. The second scenario, more applicable for mosquito control, is to release SSIMS agents as eggs. Only males will survive developmental stages in the absence of tetracycline, and thus sex-sorting occurs in the field. The latter release scenario was simulated here.

We modeled SSIMS with three independently segregating genetic loci. With allele names representing the same molecular components described in the previous section, SSIMS agents have the combined genotype ‘PPTTLL’, and wild-type have the combined genotype ‘ppttll’. As shown in Figure 2d, the SSIMS strategy is temporally self-limiting. As we simulate the release of eggs/larva there is no field amplification resulting from SSIMS males mating with SSIMS females. We observed population suppression that scaled with increased release rates. This approach is more strongly temporally self-limited than FL alone, as the genotype cannot be passed on to future generations (note difference of persistence between FL and SSIMS in Figures 2a and 2d).

To achieve moderate field amplification of the suppressive effect afforded by genetic incompatibility, we modeled a second combinatorial control strategy named Field-Amplified Male Sterility System (FAMSS). FAMSS is similar to SSIMS except that the programmable transcription activator is only expressed in germ-line tissue (Figure 2e). With this genetic design, hybrids from wild-type ^×^ engineered matings would be viable but sterile^37^. This provides one extra generation of field-amplification compared to SSIMS. Our simulations (Figure 2e) reflect this amplification, with suppressive behavior that is stronger and more long-lasting than SSIMS. Compared to FL alone, FAMSS provides a stronger suppressive effect (Supplementary Figure S2) and is more stringently temporally self-limiting, with abrupt removal of the FAMSS genotype after approximately fifty time steps (Figure 2e).

### Robustness of FAMSS and SGD to genetic resistance

Because the FAMSS strategy is self-limiting, a more realistic population suppression program would involve an initial bolus release of FAMSS individuals, followed by periodic small scale releases. These periodic releases will keep the population suppressed even in the event that new wild-type mosquitoes immigrate into the treated region. We simulated such periodic release programs for FL, SGD, SSIMS, and FAMMS with varying release numbers and frequencies (Supplementary Note 3). SGD was able to completely eradicate the wild-type population in every periodic release schedule. While this represents an unconventional and aggressive strategy for SGD release, we simulate it here for the sake of comparison and note that the decreased time to eradication will result in improved robustness to genetic resistance. Of the remaining strategies, FAMSS was the most powerful and was able to locally eradicate the wild-type population with an initial bolus release of 90% followed by weekly release of 800 eggs/larva.

Next, we modeled the stochastic emergence of genetic resistance for each control strategy (Figure 3). For SGDs to function ideally, all nuclease-induced double-strand DNA breaks would be repaired by homology-directed repair, using the drive allele as a repair template. However, if the double-strand DNA break repairs by non-homologous end joining (NHEJ) or microhomology mediated end joining (MMEJ), the repaired locus can contain mutations that confer resistance to future cutting (Figure 3a)^25,36^. We model this as the conversion of a wild-type allele ‘W’ to a resistant allele ‘R’. It is also possible that the NHEJ repair will create a non-functional allele, which we designate ‘I’.

**Figure 3.**
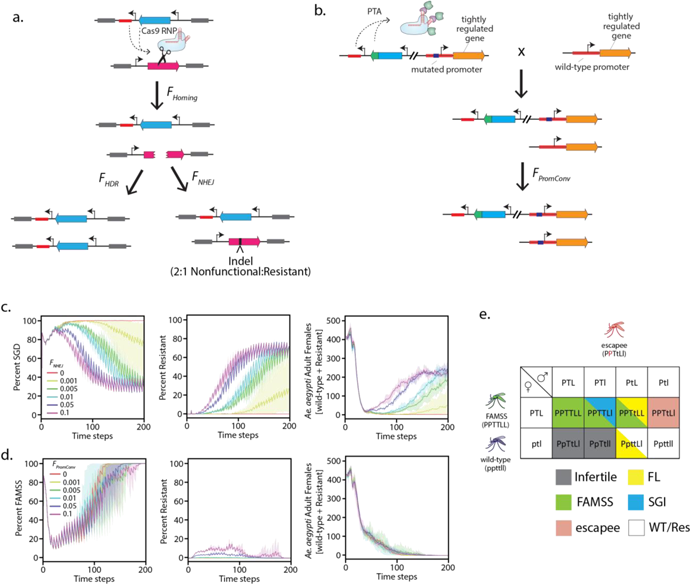
Performance of SGD and FAMSS in the context of genetic resistance. Mechanisms of simulated genetic resistance for SGD (a) and FAMSS (b) with key model parameters annotated. Performance of SGD (c) and FAMSS (d) with initial release rates of 90% GE mosquitoes and 10% wild-type, followed by release of 800 GE larvae every 5 time steps. Plots show the percentage of the total population comprising mosquitoes with the GE biocontrol genotype (left), genetically resistant mosquitoes (center), and the number of adult females with either wild-type or resistant genotypes (right). Frequency of NHEJ or promoter conversion are given in the inset of the left plot. (e) Punnett square of possible mating events between promoter conversion escapee genotype and wild-type or FAMSS mating partners.

Empirical measurements of resistance mutations in laboratory populations of *Ae. aegypti* and *D. melanogaster* point to complex and multifactorial rates of resistance that arise from differences in NHEJ repair frequency in different cell types^25,36^. To simplify this process in our model, we assume resistance arise only from NHEJ during gametogenesis. Keeping the homing frequency fixed at 98%, we modeled rates of NHEJ from 0% to 5%. In our model, an NHEJ rate of 2-2.5% closely approximates the experimentally observed behavior, with SGD alleles increasing in the population for approximately five generations before dissipating due to selection for resistant alleles (Supplementary Note 4).

Genetic resistance to SGI (and by extension to SSIMS or FAMSS) could emerge from silencing of the PTA, sequence diversity in the region targeted by the PTA, and by promoter conversion in the hybrid embryo. The first two mechanisms of resistance can be buffered by engineering strategies^18^ and were not observed frequently in a model yeast system. Promoter conversion occurs when the engineered resistant promoter of the SGI parent replaces the susceptible wild-type promoter in the hybrid embryo (Figure 3b). This form of resistance occurred at a frequency of approximately 10^−3^ in yeast^18^. In our model, we simulate resistance by promoter conversion using a by allowing offspring of FAMSS and wild-type parents (‘PPTTLL’ and ‘ppttll’, respectively) to have a viable genotype of ‘PPTtLl’ at a rate defined by the promoter conversion frequency parameter.

To determine how the FAMSS and SGD strategies perform when challenged with genetic resistance, we ran models simulating a periodic release of biocontrol agents with frequencies of promoter conversion or NHEJ, respectively, spanning three orders of magnitude (Figure 3c/d). For each rate of NHEJ tested, resistance emerged and drove down the frequency of SGD agents in the simulation (Figure 3c). Complete population suppression occurred before the emergence of resistance only at NHEJ rates of 10^−4^ or less, in agreement with mathematical modeling of a population this size^38^.

In stark contrast to the behavior of SGD, the FAMSS approach was robust to the emergence of resistance by promoter conversion (Figure 3d). Even with rates of promoter conversion of 0.1 (*e.g*. one in ten of the progeny between FAMSS and wild-type survive with a genotype of ‘PPTtLl’), the FAMSS genotype quickly went to fixation and the dynamics of population suppression mirrored the simulation in the absence of resistance. This can be explained by examining the fitness of resistant individuals in relation to wild-type or FAMSS (Figure 3e). The ‘PPTtLl’ escapee genotype is less fit than either the wild-type or FAMSS genotype. Half of the offspring between an escapee and a wild-type would be inviable as a result of inheriting a programmable transcription activator ‘T’ and susceptible promoter ‘p’. Half of the offspring between an escapee and a FAMSS mosquito would effectively regenerate the FAMSS genotype. Because the wild-type genotype continues to be suppressed in the presence of escapees, the periodic addition of more FAMSS mosquitoes gradually dilutes out the escapee genotype, leading to robust population suppression (Figure 3d).

### Behavior of FAMSS and SGD for spatially-defined population control

The ability of FAMSS or SGD strategies to rapidly suppress a population of wild-type *Ae. aegypti* would be counteracted in real-life applications by wild-type individuals immigrating into the treatment area. To examine this scenario, we performed a spatially-explicit population suppression experiment by simulating 49 individual populations on a 7 × 7 grid. All 49 cells were seeded with a steady-state starting population of wild-type mosquitoes (approximately 4000 individuals). A 3 × 3 grid in the center was treated with GE population suppression agents (Figure 4a). Each sub-population was monitored across 200 time steps (Figures 4b-d). At each time step, adult mosquitoes could stochastically emigrate out of their current cell and into one of eight neighboring cells (diagonal moves are permitted). The 7 × 7 grid was simulated on a toroid landscape, so agents in the top row could emigrate ‘up’ out of their cell and into the bottom row. We simulated the same release schedule as described above (90% initial release followed by 800 individuals every 5 time steps), with a migration rate of 0.01 for both SGD (Figure 4c) and FAMSS (Figure 4d) strategies.

**Figure 4.**
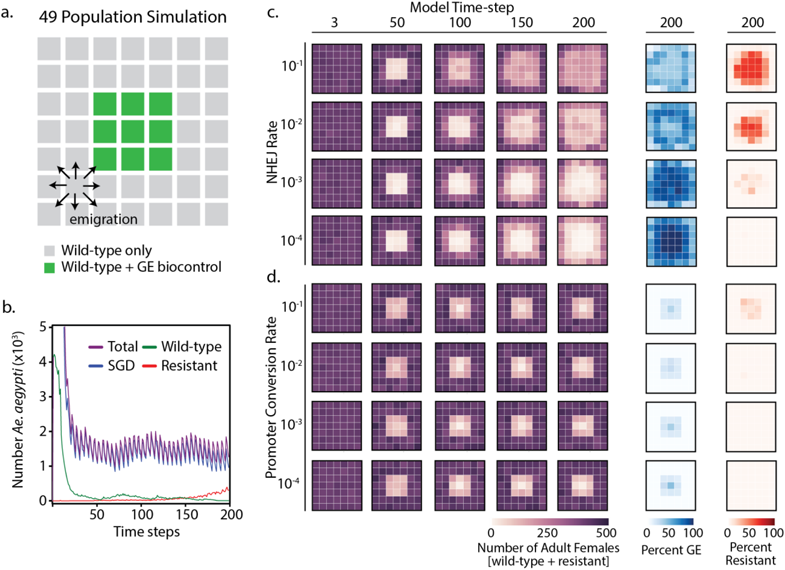
Spatially-explicit simulations of SGD and FAMSS biocontrol strategies in the context of genetic resistance. (a) Key showing location of populations treated with GE biocontrol strain. Each square represents a separate simulation model. Arrows show allowable migration routes from a single population at each time step. (b) Time-course data from a single population treated with SGD mosquitoes showing numbers of *Ae. aegypti* with genotypes represented in the inset legend. This model reflects data from the population in row 5, column 4 of the SGD simulation with *F_NHEJ_* = 10^−3^. (c) Performance of SGD in spatially explicit simulation with different rates of NHEJ (vertical axis). Plots show number of wild-type and resistant adult females (left, various time steps), and percentage of the final population comprising SGD mosquitoes (center, blue) or genetically resistant mosquitoes (right, red). (d) Performance of FAMSS in spatially explicit simulation with different rates of promoter conversion (vertical axis). Graphs show analogous data as (c), but with blue representing the final population of FAMSS mosquitoes.

Two important observations were made from the spatially-explicit treatment model using SGD agents (Figure 4c). First, the SGD genotype did not remain confined to the treatment cells. SGD individuals quickly spread throughout the 7 × 7 grid, where they transiently suppressed untreated *Ae. aegypti* populations. Second, a wave of resistant organisms shortly followed and also spread throughout the grid. This result is not surprising and has been shown by others employing different modeling techniques^38,39^. Second, SGD is less robust to emergence of genetic resistance in the multi-population simulation compared to the single population simulations. In single populations models, rates of NHEJ of 10^−3^ were sufficiently low to prevent the emergence of resistance in two of three replicates (Figure 3c) and rates of 10^−4^ saw no resistant individuals emerge (data not shown). In contrast to this, we consistently see resistant organisms emerge in the multi-population simulations with NHEJ rates of 10^−4^, signifying that SGD is less robust to challenges by genetic resistance in the multi-population simulations. Each mating event *en route* to full eradication is an opportunity for resistance to evolve and spread, and more mating events are required for suppression in the multi-population simulation. Importantly, this is not just function of increasing total population size^38^, but the rate of migration between sub-populations.

In contrast to SGD, the FAMSS approach maintained a spatially-limited zone of population suppression throughout the experiment (Figure 4d), confined to the 3 × 3 grid into which the biocontrol agents are released. Also, the robustness to genetic resistance does not change as a function of population size or structure. As is the single-population simulations, the frequency of resistance mutants in the population does not grow to fixation, even at relatively high rates of promoter conversion.

## Discussion

Our agent-based simulation model reproduces the overall behavior of several GE biocontrol strategies to suppress or replace populations of the important disease vector, *Ae. aegypti*. Embedded in the model are genetic rules that allow us to simulate female lethality, suppression gene drives, synthetic genetic incompatibility, and combination treatments. We have focused our experiments in this study to compare FAMSS with SGD in the context of genetic resistance. Performance differences between FAMSS and RIDL or Sex-ratio biasing do exist, but are more strongly impacted by parameters like fitness or mating competitiveness. Meaningful comparisons between these approaches will require empirical measurements of these parameters in engineered insects^35^.

We have incorporated into our models the likely mechanisms of genetic resistance for both FAMSS and SGD approaches. Resistance to SGD is well-documented in experimental systems ^25,36^ and has tempered the enthusiasm of using meiotic gene drives for population suppression^40^. Several approaches to slow the evolution of resistance have been proposed, such as employing multiple guide RNAs^38^ or manipulating the ratio of homology directed repair to NHEJ^41^. As the field makes progress towards overcoming resistance to SGDs, it is important to consider the performance in spatially-explicit applications. We show here that a latent population of wild-type organisms that can slowly migrate into a treatment area will exacerbate the problem of resistance, and require that rates of resistance are several orders of magnitude lower than what would be sufficient to suppress an isolated population.

Our proposed FAMSS strategy fares well when challenged with genetic resistance, both in isolated populations and in spatially-explicit models. The engineered SGI genetic construct essentially behaves as a single locus underdominant Dobzhansky-Muller Incompatibility (DMI)^42–44^, which act as stable species barriers to gene flow in neighboring populations^45^. The population genetics of DMIs are thought to be the driving force behind natural speciation, and the sheer number of species that have existed on our planet speaks to the difficulty in overcoming these barriers once they exist^43^. The FAMSS strategy we describe here, and the SGI approach in general, is robust to high frequencies of genetic resistance for the same reason.

We considered only a single mechanism of resistance, namely promoter conversion, although other mechanisms are possible. We have omitted other mechanisms of resistance because these can be addressed with simple engineering solutions. For example, silencing or mutation of the PTA can be circumvented by creating a positive selection module using an essential gene^18^. Resistance caused by underlying sequence diversity in the population would have a similar fate as promoter conversion mutants. The surviving hybrids will still one copy of the PTA and will be less fit than wild-type in subsequent mating events. Further, if two mutually incompatible FAMSS strains are engineered that drive overexpression of different target genes, these could be released in an alternating fashion. Genetic escapees from the first treatment will receive half of their genetic material from the first FAMSS strain, and can be suppressed with the second FAMSS strain. Unanticipated mechanisms of resistance will be learned from characterizing SGI insects, which are currently underway^26^.

Our spatially explicit model demonstrates the potential for FAMSS to provide long-term suppression of local mosquito populations even when challenged with genetic resistance. The migration rate used in our models (1% chance of migration per adult per time step) is reasonable given the recapture rates of mosquitoes in the field studies on which this model is based^29^. If the migration rates in a specific geographic region are greater or less that those modeled here, it will affect the numbers of mosquitoes that need to be released, but will affect the overall performance of the biocontrol strategy. The same is true of relative fitness and mating competitiveness of FAMSS versus wild-type males. As empirical measurements of these variables become available for FAMSS mosquitoes, we will update our model to improve its accuracy.

One intriguing possibility is to use FAMSS or SGI mosquitoes as a safeguard to limit the spread of a SGD release^46^. If the two genotypes were incompatible (*i.e*. the SGD strain is has a wild-type promoter at the FAMSS target locus), then a population of FAMSS mosquitoes could be pre-released to form a ‘firewall’ that will limit the spread of the SGD. For example, a resident populations of FAMSS or SGI mosquitoes could be maintained around a port-of-call with SGD released in interior of an island. By decreasing the effective population size through which an SGD could spread, FAMSS could improve the practical efficacy of SGD or other threshold-independent gene drives. Similarly, release of a large bolus of FAMSS agents could function as a fire blanket to immediately suppress an accidental or nefarious release of SGDs organisms.

In conclusion, we have described and simulated a new population control strategy that hardwires genetic incompatibility and sex-sorting into the genotype of a GE biocontrol strain. We show that this strategy is robust when challenged by high rates of genetic resistance in stark contrast to CRISPR/Cas9-based meiotic gene drives.

## Methods

### Agent-based simulation model

Our agent-based simulation model was written in Python^47^ on the Mesa platform^48^. Model parameters (of which select parameters are listed in Figure 1B) including density dependent and density independent survival rates for each life-stage, sex-ratio of new eggs, number of eggs per gonotrophic cycle, number of time steps for each life-stage, and initial number of wild-type agents were not varied during population control experiments. Variable parameters defined prior to each simulation include starting number of GE organisms, genotype of GE organisms, frequency of homing and NHEJ (SGD simulations only), frequency of promoter conversion (FAMSS/SSIMS/SGI simulations only), total number of time steps, number of GE organisms released at periodic time steps, number of time steps between periodic release, and migration rate. Agents added to the model at time step 0 are seeded at random life-stages, with 95% at life=stage 10 (last larval stage) or earlier and 5% at adult stages. Annotated Python scripts for single population simulations and 7 × 7 grid simulations as well as README instruction files are available on GitHub.

### Rules for behavior of GE control strategies

The genotype of each agent in the simulation is stored as the attribute self.genotype. To model FL, SGI, or FAMSS, each agent is given a six letter genotype, with two ‘promoter’ alleles (P or p), two PTA alleles (T or t), and two female lethality alleles (L or l). In each case, the lowercase allele designates wild-type. During the reproduction step, the genotype of each new egg produced is determined by a stochastic selection of alleles from each parent. Unless otherwise noted, each allele has a 50% chance of inheritance. Sex is determined stochastically with a 50:50 male:female sex ratio. All eggs are placed into the model scheduler and grid regardless of genotype.

To model SGD, we use a simplified two letter genotype for each agent in the model. Allele names all correspond to a single meiotic drive locus and can designate a gene drive (G), wild-type (W), resistance mutation (R), or recessive lethal mutation (I). The R and I genotypes only arise when modeling NHEJ (discussed below).

Genotype viability is assessed during the progression from lifestage 1 (egg) to lifestage 2 (first instar larva). Individuals carrying a PTA (i.e. a T allele) AND a wild-type target promoter (*i.e*. a p allele) are inviable and are removed from the model. Females carrying the female-lethal construct (*i.e*. a L allele) are inviable and are removed from the model. For SGD populations, any organism lacking at least one W allele or R allele is inviable.

### Genetic resistance in simulation model

We model promoter conversion events in SGI or FAMSS populations during the reproduction step. In any new egg that would be heterozygous at the promoter location (i.e. Pp or pP), the wild-type ‘p’ allele is changed to the resistant ‘q’ allele stochastically at a promoter conversion frequency specified at the start of the simulation. Because the rules for embryonic lethality require the co-existence of ‘T’ and ‘p’ alleles, these ‘q’ mutants remain viable. Functionally, this is equivalent to replacing the ‘p’ for an engineered ‘P’, but the ‘q’ designation facilitates future tracking of the mutated allele.

We model resistance in the SGD simulations at the stage of gametogenesis. For gene drive carriers (‘WG’ or ‘GW’), the ‘G’ allele is passed on at a frequency of:

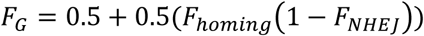

Where F_*homing*_ is the homing frequency and F_*NHEJ*_ is the frequency of non-homologous end joining. Best case gene drive simulations use a homing frequency of 100% and a non-homologous end joining frequency of 0%. The gamete will inherit the G allele from a heterozygous parent stochastically if a random number between 0 and 1 is less than or equal to *F_G_*.

Our model simulates non-homologous end joining if the random number generated, N, lies in the following range:

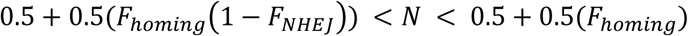

In this case, a resistant allele ‘R’ is passed on one third of the time and a haploinsufficient allele ‘I’ is passed on two thirds of the time. These probabilities are based on the random likelihood that an indel generated by NHEJ will result in an in-frame coding DNA sequence (CDS). Our model assumes that any NHEJ repair that generates an in-frame CDS will still be haplosufficient and will resist further cutting by the drive nuclease. This is likely a slight overestimate of resistance given a particular NHEJ frequency, but is reasonable for the purposes of modeling.

